# Detection of epistatic interactions with Random Forest

**DOI:** 10.1101/353193

**Authors:** Corinna Lewis Schmalohr, Jan Grossbach, Mathieu Clément-Ziza, Andreas Beyer

**Author notes:** These authors contributed equally to this work (Corinna Lewis Schmalohr, Jan Grossbach).

## Abstract

In order to elucidate the influence of genetic factors on phenotype variation, non-additive genetic interactions (i.e., epistasis) have to be taken into account. However, there is a lack of methods that can reliably detect such interactions, especially for quantitative traits. Random Forest was previously recognized as a powerful tool to identify the genetic variants that regulate trait variation, mainly due to its ability to take epistasis into account. However, although it can account for interactions, it does not specifically detect them. Therefore, we propose three approaches that extract interactions from a Random Forest by testing for specific signatures that arise from interactions, which we termed ’paired selection frequency’, ’split asymmetry’, and ’selection asymmetry’. Since they complement each other for different epistasis types, an ensemble method that combines the three approaches was also created. We evaluated our approaches on multiple simulated scenarios and two different real datasets from different *Saccharomyces cerevisiae* crosses. We compared them to the commonly used exhaustive pair-wise linear model approach, as well as several two-stage approaches, where loci are pre-selected prior to interaction testing. The Random Forest-based methods presented here generally outperformed the other methods at identifying meaningful genetic interactions both in simulated and real data. Further examination of the results for the simulated and real datasets established how interactions are extracted from the Random Forest, and explained the performance differences between the methods. Thus, the approaches presented here extend the applicability of Random Forest for the genetic mapping of biological traits.

**Author summary:** The genetic mechanisms underlying biological traits are often complex, involving the effects of multiple genetic variants. Interactions between these variants, also called epistasis, are also common. The machine learning algorithm Random Forest can be used to study genotype-phenotype relationships, by using genetic variants to predict the phenotype. One of Random Forest’s strengths is its ability to implicitly model interactions. However, Random Forest does not give any information about which predictors specifically interact, i.e. which variants are in epistasis.

Here, we developed three approaches that identify interactions in a Random Forest. We demonstrated their ability to detect genetic interactions using simulations and real data from *Saccharomyces cerevisiae*. Our Random Forest-based methods generally outperformed several other commonly used approaches at detecting epistasis.

This study contributes to the long-standing problem of extracting information about the underlying model from a Random Forest. Since Random Forest has many applications outside of genetic association, this work represents a valuable contribution to not only genotype-phenotype mapping research, but also other scientific applications where interactions between predictors in a Random Forest might be of interest.

## Introduction

Understanding the genetic components underlying biological traits has been a central focus of genetic research in humans, as well as in model organisms and agriculturally important species [1–4]. Many phenotypes, including quantitative traits (QTs) such as body height in humans, or growth rate in yeast, follow complex inheritance patterns and are influenced by a multitude of genetic variants [5,6]. Numerous genetic variants have been associated with a wide range of phenotypes in various species (e.g., through Genome-wide association studies, GWASs, and quantitative trait loci, QTL, studies). However, these variants can cumulatively usually only explain a subset of the heritability of each trait, a phenomenon that was termed ’missing heritability’ [7].

This inability to fully explain the heritable portion of phenotype variation can in part be attributed to insufficient consideration of the genetic background and interactions between genetic variants (i.e., epistasis) [8]. Epistasis describes the phenomenon when the effect of a genetic variant on a phenotype depends on the allele(s) of one or more ’modifier’ variant(s) [9]. In the context of QTs, epistasis represents non-additive interactions between loci, i.e. situations where the effect of two or more genetic variants on a QT differs from the sum of their individual (i.e., marginal) effects. Two types of epistasis can be distinguished, which are defined by their equivalent logical operations (S1 Fig). Firstly, there is AND-epistasis, where an allele at a locus enhances the effect of another locus. Secondly, XOR-epistasis describes cases where the effects of alleles at two loci are diminished when they occur together [10–12]. Epistasis is often interpreted as a functional relationship between genes, since the interactions can be explained by the loci influencing the same pathway or biological process. Identifying interacting genes can thus contribute to elucidate disease mechanisms [13,14], to identify common biological functions of genes, and to ultimately reconstruct genetic interaction networks [10,15–18].

A wide variety of methods to detect epistasis for discrete traits have been developed [19–23]. However, there is a lack of accurate methods for the detection of interactions between genomic loci that affect QTs. These are often still investigated using parametric methods, for example by evaluating pair-wise linear models (LM) for all possible locus pairs to test for non-additive interactions (for example in [24]). Such exhaustive approaches suffer from computational and statistical burdens induced by the combinatorial number of tests that have to be performed [25]. Therefore some authors use a two-stage approach where loci are pre-selected or weighed prior to interaction testing. The criteria used for the pre-selection can include the presence of marginal effects, knowledge from external databases, or data-mining approaches such as machine learning methods with variable selection [24, 26–29]. In addition, many existing methods make assumptions about the distribution of the data and the type, scale, and order of interactions, which may not apply to real data [25,30].

Several studies have shown that Random Forest (RF) outperforms other QTL mapping methods at identifying QTL and discrete trait associations [31–35]. This was mainly attributed to its ability to account for interactions [12,36]. A RF consists of an ensemble of classification and regression trees (CARTs). In each CART, the data is repeatedly split into two groups based on the predictors (in this case genetic loci) that explain the most variance of a certain outcome variable (in this case the phenotype), see also S2 Fig. Interactions are implicitly incorporated into the model through the hierarchical succession of predictors. Notably, RF is not limited in the order of interactions it can model. The predictive power and robustness of RFs stems from two separate random sampling steps [37]: (i) each CART is grown on a bootstrap sample of the data, and (ii) only a random subset of loci is considered at each split. There are several importance measures that can be used to rank predictors based on their influence on the outcome variable, which, in the context of genetic mapping, can be used to determine which loci influence the phenotype [31]. RF is model free and non-parametric in the sense that it does not assume any particular genetic model, or estimate any parameters [37].

A notorious problem of RF remains that, although it is able to take interactions into account, it does not specifically *detect* them. That is however necessary for identifying epistatic interactions between loci. The high number of studies that attempted to detect interactions in RFs or other decision tree-based methods highlights the potential of RF to capture interactions [38–45]. Some of these approaches employed pairwise importance measures, which do not provide a statistical significance level of the interactions without computationally expensive permutation-based statistics. In addition, these methods were not sufficiently tested and compared to other approaches, and/or are only applicable to discrete phenotypes. For example, we previously proposed an approach for extracting interactions from RFs [41], which relied on computationally expensive permutations and required very large RFs. This study partly builds on this previous work (i.e., the splitA approach), replacing the empirical statistics with a more powerful parametric testing approach.

We aimed at creating a method that can extract interactions between genetic loci from a RF, evaluate their significance in a permutation-free way, and is applicable to QTs. To that end, we conceived three different tests that exploit specific signatures in the RF caused by interactions. We termed these RF-based methods ’paired selection frequency’ (pairedSF), ’split asymmetry’ (splitA), and ’selection asymmetry’ (selA). Because they are expected to complement each other in different interaction scenarios, an ensemble method was also conceived by combining the *p*-values of the respective approaches. We compared these four RF-based methods to other common approaches for the detection of interactions between QTL, including an exhaustive approach that tests for linear interactions between all possible locus pairs, and several two-step approaches, where loci are pre-selected prior to interaction testing. We chose three criteria to pre-select loci: (i) an exhaustive *t*-test for marginal effects, (ii) LASSO-based feature selection, and (iii) Random Forest importance measures. To evaluate the strengths and weaknesses of the different methods, we applied them to various simulations of QTs. We then compared their performances on two real datasets comprising of expression data and growth trait data from high-throughput *S. cerevisiae* crosses. For the real data, we measured the method’s performance based on their ability to recover previously published genetic interactions identified through double gene knock-outs [18].

## Results and discussion

### Description of selected methods

The three RF-based methods each investigate specific signatures of interaction in a RF. Detailed descriptions of each method are given in the Methods section. In short, the pairedSF investigates whether two loci are more likely to be selected in the same CART than would be expected by chance (based on their individual selection frequencies). The second approach, splitA, examines the difference in the effect of a locus on trait value distributions, depending on the allele of another, potentially interacting locus used earlier in the same path of a CART. Finally, the selA approach exploits the fact that loci in AND epistasis have different probabilities of being selected for a split in the forest depending on a previous partitioning on the other interacting locus. Finally, we combined the respective *p*-values of these three approaches to obtain an ensemble method.

We chose several commonly used approaches to compare to our RF-based methods. First, we applied an exhaustive linear model (LM) approach, where the significance of the interaction term based on an ANOVA was used to test for interactions between all possible locus pairs. However, because our four RF-based methods benefit from the feature selection properties of the RF, a comparison to only an exhaustive approach would be inadequate. Therefore, in order to evaluate to what extent our methods’ performances depended on this feature selection of RF, we included three two-step approaches in the analyses. In these methods, loci were pre-selected prior to interaction testing using the LM procedure. First of all, we implemented a ’*t*-Test 2step’ approach, where loci were pre-selected based on whether they manifest marginal effects according to an exhaustive *t*-test screen. In addition, we used the variable selection properties of the LASSO to pre-select loci based on their coefficients (’LASSO 2step’), as suggested previously [46, 47]. And finally, we applied a ’RF 2step’ approach, in which we exploited the feature selection properties of the RF algorithm to pre-select loci based on their importance values, as previously suggested [48–52]. The latter differs from our suggested RF-based epistasis detection methods in that the RF is only used to pre-select the loci, but the RF structure is not exploited to identify the interactions. Therefore, the genetic background modeled by the RF is not taken into account for the actual interaction testing with the RF 2-step approach. For each two-step approach, we both tested for interactions *within* preselected loci (’both’ loci have a marginal effect), and between the pre-selected loci and all others (only ’one’ locus with marginal effect), as previously proposed [53]. So in total, there were the four RF-based methods (the three individual approaches and their ensemble), an exhaustive approach, and a total of six two-step approaches (three pre-selection criteria, with the ’both’ and ’one’ strategy each).

### Simulated data

Based on genotypes of a widely used yeast recombinant panel [54], we simulated phenotypes using 19 different scenarios of different combinations of marginal and epistasis effects with differing effect sizes, different interaction types (two-way and three-way AND epistasis, and XOR epistasis), and varying levels of noise. Since each scenario was simulated 32 times, we evaluated the performance of each method based on the proportion of simulations where the *p*-values of the truly interacting loci ranked among the smallest 5% of *p*-values. It should be noted that the way the interactions were modeled (i.e., using a formula with marginal and interaction terms) conforms to the model used for a LM, which might favor the LM-based approaches (i.e. the exhaustive LM and the two-step approaches). The performances of the different methods on four representative scenarios are shown in Fig 1A-D, and are summarized in Fig 1E.

**Fig 1.**
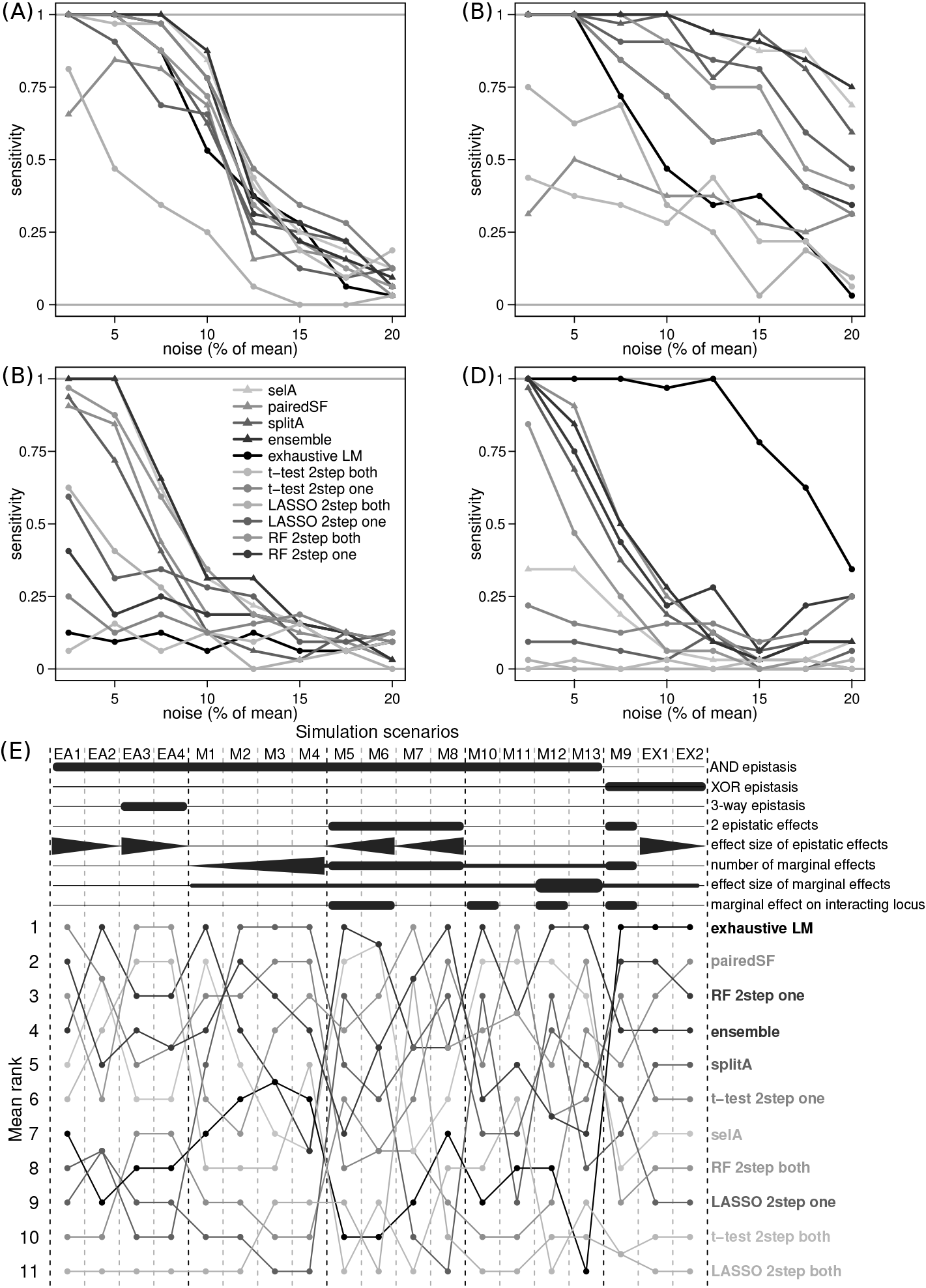
Sensitivity of methods based on simulations. Sensitivity was measured by the proportion of simulations where the *p*-value of the truly interacting locus pair was below the lower 0.5-percentile of *p*-values (i.e. 99.5% of *p*-values were higher). Panels (A) to (D) show the results for four representative simulation scenarios. (A) Pure (no marginal effects) two-way AND epistasis, scenario EA1; (B) Two-way epistasis with an additional marginal effect on one of the interacting loci (scenario M12); (C) Two-way epistasis with an additional marginal effect on a distinct locus (scenario M13); (D) Pure two-way XOR epistasis (scenario EX1). (E) Summary of the results for all simulation scenarios. The mean rank over all noise levels is depicted for each method and scenario. The features of each simulation scenario are indicated above the upper panel.

In general, all RF-based methods were able to detect pure AND epistasis (Fig 1A and S3 FigA). Among them, the ensemble method generally performed best, and it also slightly outperformed the other methods at recovering simulated interactions (Fig 1E). The RF-based approaches profited a lot from cases where at least one of the interacting loci had a marginal effect (Fig 1B, S4 and S5 Figs). This represents one of the limitations of RF: it relies on the presence of marginal effects for the selection of loci in the CARTs. Correspondingly, the two-step approaches applied here also rely on the presence of marginal effects on at least one of the interacting loci. However, AND epistasis will, even in the absence of underlying true marginal effects, lead to a phenotype difference for the interacting alleles separately (i.e., a ’quasi-marginal’ effect), promoting their selection in the RF. For example, considering a case of pure AND epistasis, where the phenotype is only affected if both loci have an altered allele (S1 FigB), each locus taken alone will appear to have a marginal effect on the phenotype. This is the reason why the RF-based methods were still able to uncover the interactions in scenarios without any marginal effects (Fig 1A). In addition, interacting loci without any marginal effects are unlikely in a real biological setting [55], which alleviates the severity of this limitation of the RF-based approaches.

The RF-based approaches still identified the correct interactions in the presence of additional marginal effects on non-interacting loci (i.e., Fig 1C). Surprisingly, the RF-based methods were not as robust as the other methods against increasing numbers of such ’interfering’ marginal effects (S6 Fig). It should be noted, however, that the two-step RF approach, together with the two-step LASSO seemed to be most suited to identify and pre-select the truly interacting loci. This further emphasizes the advantage of multivariate approaches for modeling complex genetic regulation.

The ’quasi-marginal’ effects mentioned above induced by AND-epistasis do not necessarily arise for XOR epistasis (S1 FigC, loci A and B will not appear to have a marginal effect if examined individually), making them potentially hard to capture with RF. For our simulations, we chose a scenario that does not lead to any ’quasi-marginal’ effects. As expected, the exhaustive LM approach outperformed all other methods at the detection of this interaction type (Fig 1D and S7 FigA). However, it should be noted that aside from the exhaustive LM approach, all RF-based methods (i.e., our novel approaches and the two-step RF approach) performed better than the others. Surprisingly, adding marginal effects to both interacting loci did not facilitate the detection of XOR epistasis for any of the methods (S7 FigB). Although there are theoretically possible mechanisms that can lead to XOR epistasis for both discrete and quantitative traits, the biological relevance of XOR epistasis has been a matter of controversy [56–58]. Notably, a thorough search of the literature did not uncover any real biological examples of XOR epistasis for a quantitative trait. This lack of discoveries, however, might also be due to the fact that this type of interaction is particularly difficult to detect [58,59].

RF is claimed to be able to model higher-order interactions. Therefore we also created two simulation scenarios with three-way AND epistasis. For simplicity, we considered three-way interactions as recovered if all pairwise combinations of the three interacting loci were detected. The performance of all methods was relatively poor, however employing RF (e.g., in the ensemble method, or the two-step RF method) seemed to offer an advantage at capturing these higher-order interactions, as did the employment of the relatively sensitive *t*-test-based pre-selection (S3 FigB-C).

As expected, the three RF-based approaches complemented each other for different interaction scenarios. The pairedSF method, for example, which generally performed weaker on scenarios involving AND-type epistasis, performed comparatively better for XOR epistasis. Indeed, the ensemble generally outperformed each individual RF-based approach.

### Benchmark on expression data

Having demonstrated that our methods are theoretically able to detect epistasis on simulated data, we evaluated them on real data to test the applicability of the methods in a real biological setting. Mapping expression QTL (eQTL) is a key aspect towards understanding the relationship between genetics and phenotype. Therefore, an eQTL dataset derived from a yeast cross with 112 segregants (RM×BY) was selected [54]. The methods were applied to detect interactions between loci influencing the expression of each transcript. We evaluated the performances of the different methods by the ability to recover epistatic interactions detected in double knock-out (DKO) experiments [18], as previously proposed [41]. Since these DKO interactions were based on growth phenotypes, we limited the epistasis detection to 1,050 transcripts corresponding to essential genes (i.e., genes that are lethal when knocked-out), assuming that these were more likely to affect growth than non-essential genes. An eQTL interaction was assumed to be a true positive if genes close to the interacting loci were also found to interact in the DKO dataset [18]. We used these true positive labels to compute precision-recall (PR) curves and the area under the receiver operating characteristic (AUROC). It should be noted that the eQTL dataset differs widely from the DKO data in the type of phenotypes and genotypes studied (expression versus growth, and natural genetic variation versus whole gene knock-outs, respectively), as well as in the study design. Due to the likely small overlap between the two studies, false positive rates and false negative rates are likely grossly overestimated. Hence, an absolute performance quantification of the methods is not possible. This benchmark based on the DKO data should only be considered as the evaluation of *relative* performance differences between the methods, as measured by the enrichment of true positive calls among the detected interactions.

The PR curves of most methods and the AUROCs of all methods were better than expected by chance (Fig 2), which confirms that the DKO data is—at least to some extent—suitable as a reference for quantifying true epistatic interactions, despite the differences outlined above [41]. The RF-based methods, especially the ensemble approach, exhibited the best performance according to both the PR curves and the AUROC (Fig 2). To better understand the higher performance of the RF-based methods, we investigated the interactions for each transcript that were exclusively detected by each method (but not detected by the other approaches) and which were also considered to be true positives according to the DKO dataset (methods). For this analysis we focused on a subset of methods, i.e. the three RF-based approaches, the ’exhaustive LM’, the ’t-Test 2step one’, the ’LASSO 2step one’, and the ’RF 2-step one’. As a reference, we also extracted ’consensus’ interactions, i.e. interactions that were found by at least five of the seven considered methods, and which corresponded to interactions also found in the DKO dataset. First of all, we examined the type of interaction (i.e., AND or XOR epistasis) of the respective method-specific interactions. The pairedSF approach had the highest AUROC among the RF-based methods and primarily detected AND-type interactions (S8 Fig A). This was a somewhat surprising result, since pairedSF had a high sensitivity for XOR epistasis in the simulations. In contrast, the interactions detected exclusively by the exhaustive approach were mostly XOR interactions. This is in line with the simulations, where the exhaustive LM had outperformed all other methods for the detection of XOR epistasis. Notably, about two thirds (61%) of the ’consensus’ interactions were XOR interactions. This could be due to several reasons: (i) the consensus interactions are enriched in interactions from the exhaustive LM, simply because it was applied to the most locus pairs; (ii) transcripts in this cross are actually affected by more than 50% XOR interactions; or (iii), the interactions from the DKO dataset are enriched for XOR interactions.

**Fig 2.**
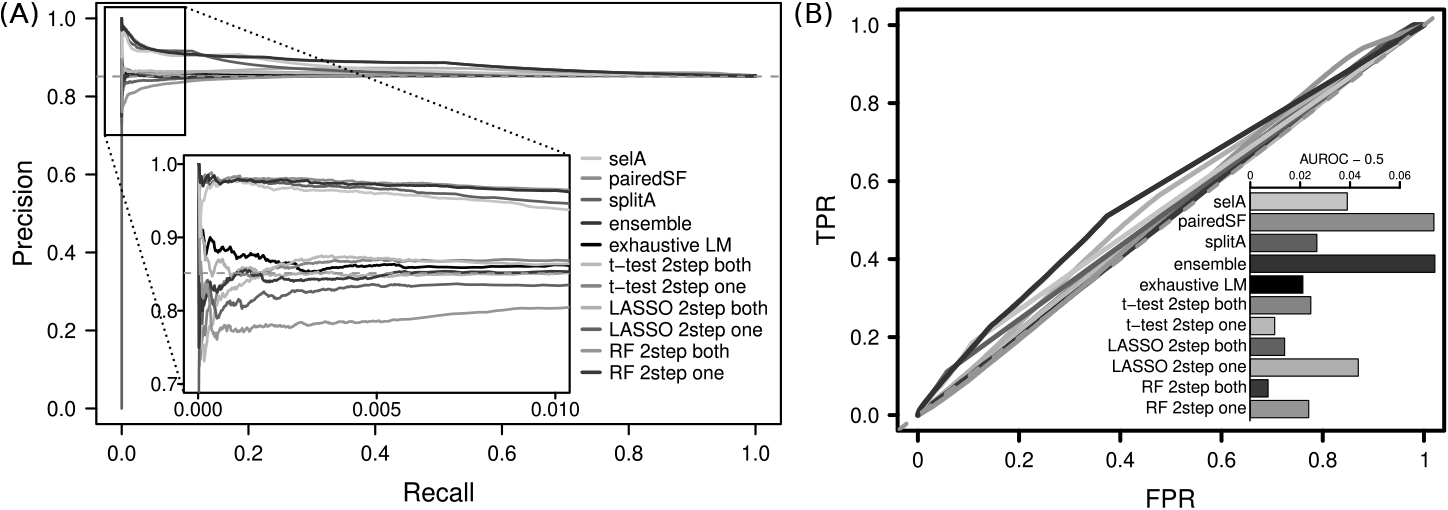
Performance on expression data. Performance was evaluated by the ability of the evaluated methods to correctly classify interacting and non-interacting genes, using a double knock-out dataset as a reference [18]. (A) Precision recall (PR) curve. A magnification of the beginning of the curve is shown in the inset. The dashed line indicates the expectation under random assignment. (B) Area under receiver operating characteristic (AUROC). The area expected under random assignment (0.5) was subtracted.

Next, we investigated whether the loci of the method-specific interactions had marginal effects (according to a *t*-test *p*-value below 0.05, without multiple testing correction) (S8 FigB). The exhaustive LM was not dependent on marginal effects. The RF 2-step and the LASSO 2-step detected 88% and 62% of the interactions, for which none of the interacting loci had a detectable marginal effect, respectively. This stresses the advantage of multivariate methods, which can unravel marginal effects that only come to light if multiple genetic variants are considered together, but not when they are tested individually, as for example with a *t*-test [25,32,33]. Notably, pure XOR epistasis is one example of an interaction that would not lead to any detectable marginal effects. The RF-based methods primarily detected interactions with additional marginal effects. This is in line with the simulations, where the RF-based methods were better at detecting interactions with additional marginal effects on the individual loci. The pairedSF was the only one of the RF-based methods that also detected interactions without marginal effects, which is one possible reason for its slightly higher AUROC compared to the other RF-based methods.

Next, we investigated whether the locus with the highest marginal effect for the respective transcript (i.e. smallest *t*-test *p*-value) was one of the method-specific interacting loci. In other words, we checked whether there was a large marginal effect present aside from the interacting loci (S8 Fig B). Contrary to expectations, the RF-based approaches did not detect more interactions outside the major marginal effects compared to the other methods.

Finally, it should be noted that the number of locus pairs tested per transcript varied a lot between the different approaches. As expected simply due to the nature of the tests, the exhaustive approach always considered all locus combinations, while the RF-based and two-step approaches tested fewer locus pairs due to their heuristic properties (S8 Fig D). The RF-based methods were more precise than for instance the two-step approaches, which investigated similar numbers of locus pairs. Therefore, the limitation that the loci have to occur together in the tree represents a valuable addition for the interaction testing as opposed just just looking at their individual importance in a tree (i.e., RF 2-step approach). The pairedSF is less restrictive about which locus pairs can be tested (i.e., it does not require the loci to occur in the same CARTs that many times and is therefore applicable more often). The splitA and selA approaches have a higher true positive rate at low false positive rates (beginning of the ROC curve) compared to the pairedSF. Thus, the few interactions detected by the splitA and selA are relatively precise, but the pairedSF has a higher recall, simply because it is applicable to more locus pairs.

A possible reason for the higher performance of the RF-based approaches compared to the other approaches could be that the RF-based methods take potential complex relationships between the genetic factors into account while testing for pair-wise interactions, whereas a LM ignores higher-order or other more complex interactions.

### Benchmark on growth trait data

For a second real data benchmark, a dataset from a distinct yeast cross with 720 segregants was selected, which encompassed measurements for several growth traits [60]. These growth phenotypes represent a more complex phenotype than gene expression, offering more opportunities for complex interactions [61]. Again, the results of the different methods were evaluated based on the recovery of interactions from the DKO dataset [18]. We focused on the phenotype ’fitness on agar’, because it was the phenotype that most closely resembled the conditions of the DKO study, so that the results for this phenotype are expected to show a higher overlap with the DKO interactions. The performances of all methods were very similar to what would be expected from random assignment, with only small differences between methods (Fig 3. None of the methods detected many interactions that were consistent with the DKO dataset. The small differences between the methods could be due to the smaller number of considered interactions: as opposed to the expression dataset, where we studied 1050 traits and 1275 loci, we now only considered one trait and 2827 loci. However, despite the small absolute number of consistent interactions detected, most epistasis detection methods found more interactions that were consistent with the DKO dataset than expected by chance. In order to test if the small improvement over random for the ’fitness on agar’ trait is relevant, we compared those results to results for the phenotype ’resistance to cantharidin’. It has been shown that the trait ’resistance to cantharidin’ is monogenic in this cross [60]; hence it is not expected to be subject to epistasis, and can therefore be treated as a negative control. Indeed, the AUROC and PR measures for the trait ’cantharidin resistance’ were close to random (S9 Fig), suggesting that the small improvement over random for ’fitness on agar’ can be interpreted as a true signal. The ensemble method showed the best performance according to PR and AUROC, together with the ’t-Test 2step one’ and ’RF 2step one’ approach. Thus, this benchmark represents supportive evidence for the applicability of the RF-based methods for real data.

**Fig 3.**
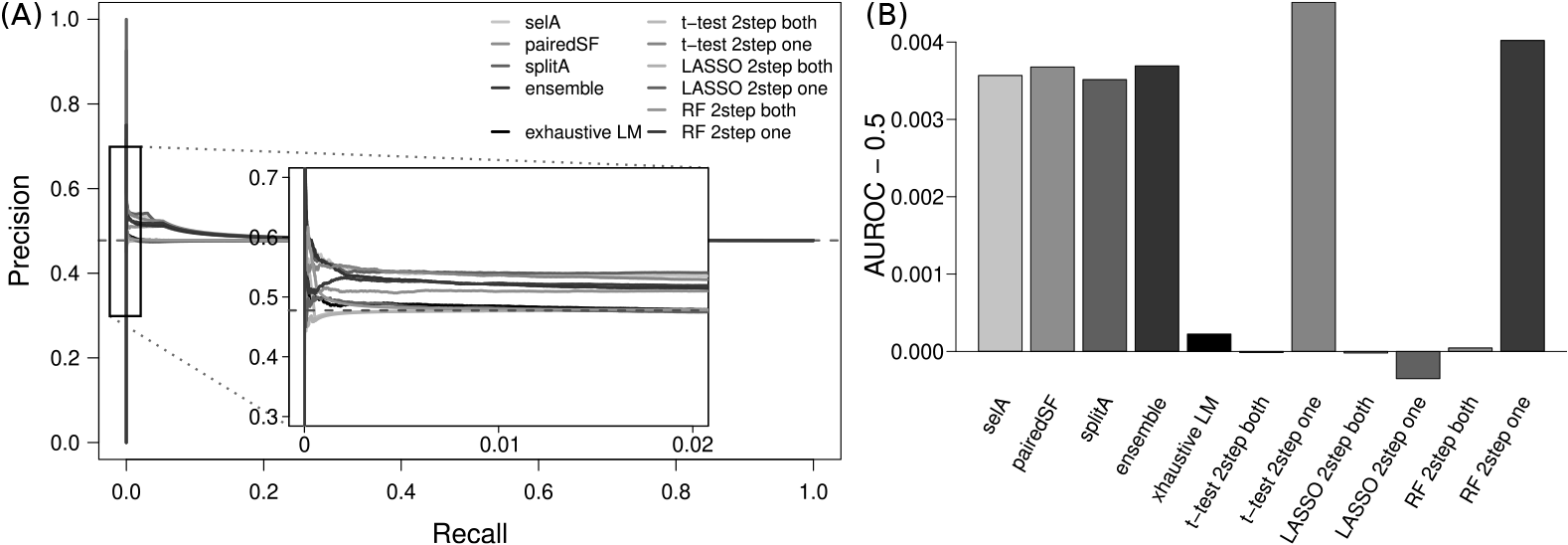
Performance on growth trait data. Performance was evaluated by the ability to correctly classify interacting and non-interacting genes, using a double knock-out dataset as a reference [18]. (A) precision-recall (PR) curve. The dashed line indicates the expectation under random assignment. (B) Area under receiver operating characteristic (AUROC), with the values expected under random assignment subtracted.

## Conclusion and outlook

We have developed three approaches that are able to extract information about interactions from a RF model. We used them to identify epistasis between genetic loci. Traditionally, simulated QTL data have been used to evaluate the performance of epistasis detection methods. Obviously, simulated data have the advantage that the true result is known, which greatly helps understanding the scenarios under which a particular method performs better or worse than another method. However, simulations suffer from the problem that they usually do not capture the full complexity of real data. Hence, we aimed to also develop a scheme for evaluating QTL epistasis detection methods on real QTL data even if a ground truth is unknown. Our study revealed that indeed the comparison with DKO data is a reasonable way for quantifying the relative performance of epistasis detection methods.

Benchmarks on simulated and real yeast cross data indicated that the RF-based methods outperform other commonly used approaches at detecting epistasis. Particularly the ensemble approach, which combines all three RF-based approaches, performed relatively well at this task. It generally achieved the highest precision for the simulated data. When applied to real data and compared to the other tested methods, it seemed to produce the biologically most meaningful results, as indicated by the recovery of high-confidence interactions that were identified in an independent study from double gene knock-outs. Therefore, the approaches for the identification of interactions within the forest introduced here represent a valuable enhancement of the applicability of RF for genetic association analyses.

Higher-order interactions (three-way, four-way, etc.) are likely to play a role for many complex phenotypes [23,62]. However, the search space for interactions grows exponentially with the order of interactions that are investigated, making an exhaustive search for three-way interactions currently infeasible. One of the advantages of RF is its ability to account for such higher-order interactions, even if they are not explicitly detectable. The feature selection properties of RF alleviate some of the multiple testing burden; hence, future work should explore the possibility to detect higher-order interactions with RF.

While studies on yeast represent a valuable resource for the investigation of the influence of genetic factors on complex traits [55], it remains a relatively simple genetic model organism, especially since it is haploid. In order for the approaches presented here to be applicable to more complex species, including humans, they have to be adapted to incorporate diploid interaction models and take dominance effects into account. Efficient implementations of RF that are applicable on a genome-wide scale in humans have already been developed [63] and RF was previously used for GWAS in humans [33,35,64]. Especially the pairedSF approach would be straightforward to extend to other species as it is not affected by dominance effects and the more complex interaction models that apply to diploid species. The efficacy of the here presented RF-based interaction detection approaches in other species warrants further investigation.

Finally, RF is a popular tool for regression and classification in various scientific applications aside from the genetic mapping problem. Therefore, the approaches to use RF for the detection of interactions presented here represent a valuable resource for various scientific applications where information about the relationship between predictors in a RF is of interest.

## Materials and methods

### Random Forest methods

All methods were implemented with custom scripts for the R statistical environment [65]. For the detection of epistasis using RF, a modification of the R implementation ’randomForest’ [66] was used. It solely differs from the original implementation by its ability to parallelize growing the trees, and to return OOB predictions also for internal tree nodes. Unless stated otherwise, the RF was always built in the same way: 30,000 trees were grown, the minimum final node size was set to 5, and all other parameters were left at the default.

RF cannot deal with missing data in the predictors (i.e., the genotypes). Therefore, missing genotypes were imputed in 100 iterations by randomly selecting the missing alleles according to the respective allele frequencies at each locus. 300 trees were grown for each imputation, and the resulting collection of forests was then united into one big forest of 30,000 trees. For the two real datasets, population structure representatives were included as covariates in the model (i.e., as predictors used to build the RF), as proposed previously [67]. For all methods, locus pairs in strong linkage disequilibrium (LD), as indicated by an absolute Pearson correlation value above 0.9, were excluded (i.e., the *p*-value was set to not available, NA).

### Random Forest paired selection frequency

This approach is based on the expectation that interacting loci are more likely to be selected in the same CART than non-interacting loci. For example, given two hypothetical epistatic loci A and B, locus B will have a higher probability to be selected for a split if the data was previously split on locus A, especially if they interact through XOR epistasis. We built a contingency table with the number of trees in which both loci were selected (number of co-occurrences *N*_*AB*_), the number of trees where the loci were selected independently of each other (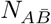 and *N*_*ĀB*_), and the number of trees none of them occurred 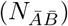. A one-sided Fisher’s exact test was applied to see if *N*_*AB*_ was bigger than expected, indicating co-dependencies between the loci. Two simplifications were necessary for this approach. For one, trees where locus combinations are used several times in the same tree were counted as just one co-occurrence. In addition, this method disregards whether two loci actually occur in the same path of a CART or not.

### Random Forest split asymmetry

Given two loci A and B in the same path of a CART (A before B), the difference in mean phenotype observed after a split on locus B depends on the result of the partitioning on locus A, as indicated by the green slopes *S*_*ABl*_ and *SA*_*ABr*_ in Fig 4. This phenomenon can be observed for both AND-type and XOR-type epistasis. This specific splitting behavior, termed split asymmetry (splitA), was already previously exploited to detect epistatic interactions [41]. Here we substantially improved the efficiency and statistical power of the approach in two ways: first, we not only consider locus pairs that occur in direct succession of each other in a CART, but also when there are splits on other loci in between them, which ultimately allows the inclusion of more observations at the same RF size. In addition, instead of using computationally expensive permutations to compute empirical *p*-values, a Student’s *t*-test is used to compare slopes.

**Fig 4.**
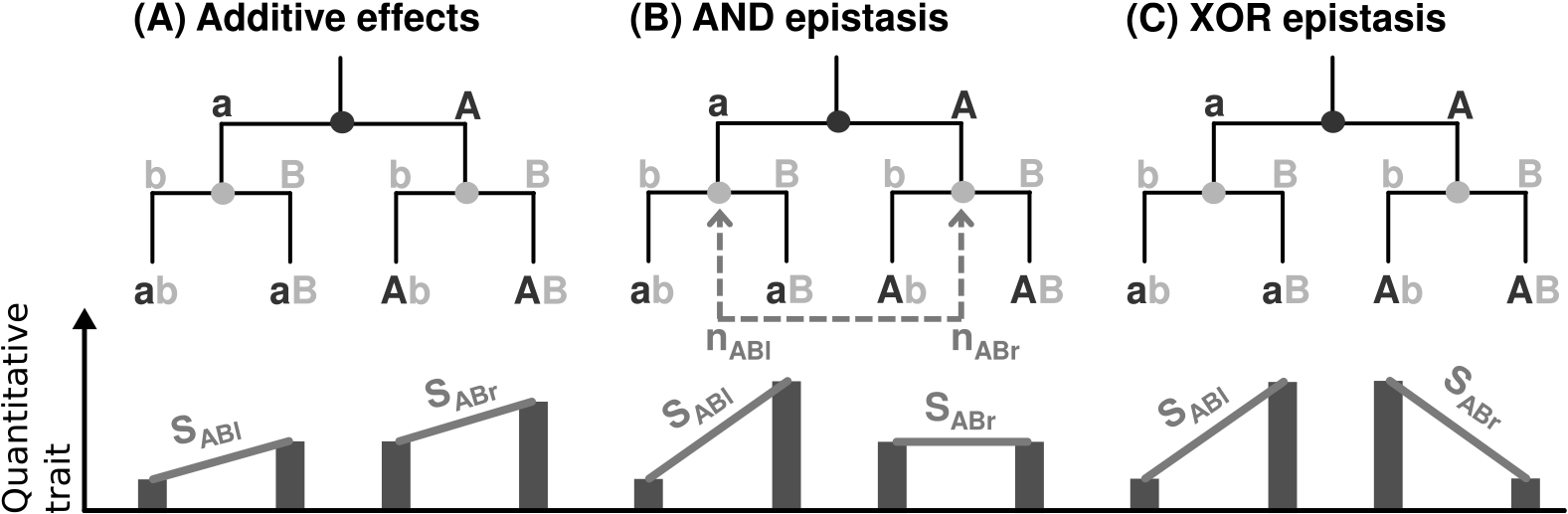
Schematic representation of the detection of epistasis from Random Forest. Shown are hypothetical subtrees that depict the splitting of data on two loci A and B that (A) do not interact, (B) are in AND epistasis, or (C) are in XOR epistasis. The latter two lead to asymmetries in the trait value distribution (indicated by the green slopes), which are exploited in the splitA approach. In addition, there are unequal probabilities for the selection of locus B for the two partitions created by the split on A (indicated by the red dashed arrow), which is tested for in the selA approach. In practice, not only cases where two loci are used directly after each other are considered, but also cases where there are other splits between them.

To that end, we calculated the slopes by taking the difference between the out-of-bag phenotype values of the respective tree nodes for each locus pair. The sums of slopes ∑*S*_*ABl*_ and ∑*S*_*ABr*_, the sums of squares of slopes ∑*S*_*ABl*_^2^ and ∑*S*_*ABr*_^2^, and the numbers of slopes *n*_*ABl*_ and *n*_*ABr*_ (here for the case where B was used after A) are sequentially updated for the left (l) and right (r) slopes, while iterating through the CARTs in a RF. These are used to compute means (*m*_*ABl*_ and *m*_*ABr*_) and variances (*υ*_*ABl*_ and *υ*_*ABr*_) of the slopes. The *t*-statistic is then calculated as

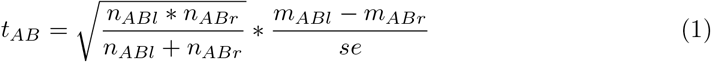

with

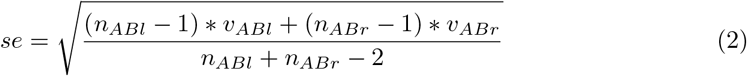

and compared to a *t*-distribution with *n*_*ABl*_ + *n*_*ABr*_ − 2 degrees of freedom. The Student’s *t*-test could only be applied if there were at least five slopes *S*_*ABl*_ and *S*_*ABr*_ each. The same test is applied to cases where locus A was used after locus B (i.e., compare *S*_*BAl*_ to *S*_*BAr*_), so that each locus pair is tested twice in two independent tests. The two *p*-values are then combined using the Fisher method [68]. If only one *p*-value could be computed (e.g., the locus pair occurred in the same trees in the order AB, but never in the order BA), only this *p*-value is used.

### Random Forest selection asymmetry

When two loci A and B interact through AND-epistasis, there is an imbalance in the frequency of splits using B after A (Fig 4, red dashed arrow). More specifically, assuming that locus B only has an effect on the phenotype given the allele ’*A*’ at locus A, but not given the allele ’*a*’, B will only be selected for a split on one side of a previous split on A. This means that there might not be enough slopes to apply the splitA approach. However, we exploit this selection asymmetry (selA) to detect epistasis by counting, for each locus pair (A and B), the number of times B is used on the left side, or on the right side after a split on A, respectively (*n*_*ABl*_ and *n*_*ABr*_).

Correspondingly, the number of splits in the tree where locus B is *not* used after a split on locus A (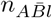 and 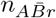) are counted (i.e., splits where any other locus than B was used after A). A binomial test of equal probabilities (without Yates’ continuity correction) is then used to test for the interdependence in the loci selection frequency, treating the total number of splits following a split on each side of A (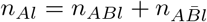 and 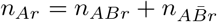) as the numbers of trials, and *n*_*ABl*_ and *n*_*ABr*_ as the numbers of successes. Locus pairs were only tested if B was used in at least five splits on either side of A. Again, each locus pair is tested twice: once for the cases where locus B is used after locus A in the tree, and also for the opposite case. The *p*-values for the two possible pairs (AB and BA) are combined using the Fisher method.

### Random Forest ensemble approach

The three RF-based epistasis detection methods described above complement each other for different types of epistasis (i.e., selA theoretically only works for AND-epistasis, and pairedSF is most likely to be able to detect XOR-epistasis). In addition, the methods are applicable in different scenarios due to different statistical requirements: splitA and selA can only be used when a locus was used at least five times on both sides, or on either side of another locus, respectively. Thus, the *p*-values generated by the splitA, selA, and pairedSF approaches were combined using Fisher’s method to create an ensemble score. Only the *p*-values of the methods that were available for each locus pair were used (e.g., if only two out of three methods were applicable for a locus pair, only the *p*-values of these two were combined).

### Exhaustive linear models

For all locus pairs (e.g., A and B) with an absolute Pearson correlation coefficient |*r*| ≤ 0.9, a linear model (LM) that models the phenotype *y* was computed according to the formula

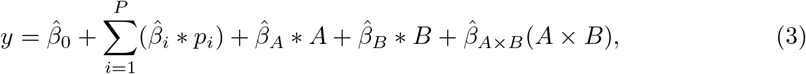

where *P* represents the collection of population structure covariates *p*_*i*_. The significance of the interaction term 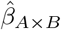 according to an F-test was then used as the significance of the locus-locus interaction. The same LM was also used for the two-step approaches to test for interactions for the pre-selected loci.

### *t*-Test-based two-step approach

This approach employs a Student’s *t*-test to preselect loci prior interaction testing. First, an exhaustive Student’s *t*-test was used to rank loci by their marginal effects. Samples with an unknown genotype for a locus were not used. It should be noted that in this first pre-selection step, population structure could not be taken into account. However, the consideration of population structure in the subsequent steps avoids the selection of false-positive effects due to population structure. The top 100 loci, ranked according to their respective *p*-values, were considered to have a marginal effect. A LM was subsequently used to test for interactions for locus pairs where either both loci had a marginal effect, or only one locus had a marginal effect.

### LASSO-based two-step approach

The machine learning method LASSO was used to preselect loci for interaction testing. Missing genotypes were handled similar to the procedure used for RF: 500 distinct imputations based on allele frequencies were created. For each imputation, a LASSO model was computed, where the phenotype was regressed against the collection of loci and population structure covariates. The LASSO algorithm leads to a sparse model by sequentially shrinking coefficients to zero according to the sparsity parameter λ. The optimal λ was determined via cross validation, using the ’cv.glmnet’ function from the ’glmnet’ R package. The loci were ranked based on the number of LASSO iterations where their coefficients were non-zero, and the top 100 were used for subsequent interaction testing using the LM procedure described above.

### Random Forest-based two-step approach

To evaluate how much our methods profited from the feature selection properties of RF, we applied a two-step approach based on RF importance values. A forest with 20,000 trees was computed. In the case of missing genotypes, they were imputed in 100 iterations as described above, and 200 trees were grown on each imputation. The commonly used importance scores ’permutation importance’ (PI) and ’increase in mean squared error’ (RSS) of each locus were extracted. These two measures were summarized in a combined score 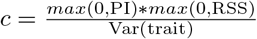. The loci were ranked based on this combined score and the top 100 were used for interaction testing.

### Performance evaluation on simulated data

We simulated trait values based on the genotypes from the widely used data from a cross between the two *S. cerevisiae* strains RM11-1a and BY4716 [54]. A set of 1275 polymorphic loci remained after collapsing identical genetic variants. Missing alleles were set to the genotype of the RM strain. In total, 19 different combinations of marginal and epistatic effects with varying effect sizes and different types and orders of epistasis were simulated according to the formula

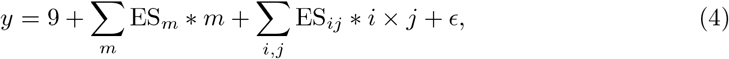

where *y* represents the phenotype values, ES_*m*_ and ES_*ij*_ represent the effect sizes for the loci with marginal effect *m*, and interacting loci *i* and *j*, respectively. We selected the baseline of 9 randomly. Normally distributed noise *ϵ* was added at eight noise levels ranging from 2,5 to 20 % of the mean. The parameters for all scenarios are listed in Table 1.

**Table 1.** Outline of the features of the different simulation scenarios.

| Scenario name | Marginal effects |  | Interaction effects |  |  |  | Overlap |
| --- | --- | --- | --- | --- | --- | --- | --- |
| | Number of marginal effects | Marginal effect size ($ES_m$) | Number of epistatic effects | Type of epistasis | Order of interaction | Epistasis effect size ($ES_{ij}$) | Marginal effects on interacting loci? ($m \in \{i, j\}$?) |
| EA1 | 0 | n.a. | 1 | AND | 2 | 1 | n.a. |
| EA2 | 0 | n.a. | 1 | AND | 2 | 0.5 | n.a. |
| EA3 | 0 | n.a. | 1 | AND | 3 | 1 | n.a. |
| EA4 | 0 | n.a. | 1 | AND | 3 | 0.5 | n.a. |
| EX1 | 0 | n.a. | 1 | XOR | 2 | 1 | n.a. |
| EX2 | 0 | n.a. | 1 | XOR | 2 | 0.5 | n.a. |
| M1 | 1 | 1 | 1 | AND | 2 | 1 | No |
| M2 | 2 | 1 | 1 | AND | 2 | 1 | No |
| M3 | 3 | 1 | 1 | AND | 2 | 1 | No |
| M4 | 4 | 1 | 1 | AND | 2 | 1 | No |
| M5 | 2 | 1 | 2 | AND | 2 | 0.5 | Yes |
| M6 | 2 | 1 | 2 | AND | 2 | 1 | Yes |
| M7 | 2 | 1 | 2 | AND | 2 | 0.5 | No |
| M8 | 2 | 1 | 2 | AND | 2 | 1 | No |
| M9 | 2 | 1 | 2 | XOR | 2 | 1 | Both |
| M10 | 1 | 1 | 1 | AND | 2 | 1 | Yes |
| M11 | 1 | 1 | 1 | AND | 2 | 1 | No |
| M12 | 1 | 3 | 1 | AND | 2 | 1 | Yes |
| M13 | 1 | 3 | 1 | AND | 2 | 1 | No |

n.a.: not applicable

Each scenario was simulated 32 times. Accordingly, the performance of each method was evaluated by counting the number of simulations where the *p*-value(s) of the truly interacting locus pair(s) was below the lower 0.5 percentile of *p*-values (i.e., 99.5% of *p*-values were higher), at each noise level. For scenarios with more than one epistatic locus pair, all of them had to be detected in order for the simulation to be considered recovered. For three-way interactions, all pair-wise combinations of the interacting loci had to be detected.

### Performance evaluation on real data using DKO data

In order to evaluate the methods’ performance at identifying interactions for real data, we used a DKO dataset as a reference [18]. This dataset consists of gene-gene interactions that were identified via co-knockouts of yeast genes using the synthetic genetic array method. In short, they tested whether double knock-outs showed a significant deviation in fitness compared to the expected multiplicative effect of the respective two single knock-outs. For this study, the results for essential (both damp-and temperature sensitive alleles) and non-essential genes were combined. Interactions which passed the lenient threshold used in the original paper (*p*–value ≤ 0.05) in at least one assay were considered interacting. We excluded pairs with genes that were removed or marked as ’dubious’ in the latest reference annotation (ENSEMBL, version R64-1-1). Overall, about 13% of all possible gene pairs were found to interact. Because QTL analysis identifies interacting loci and not interacting genes, genes had to be assigned to nearby loci (see also below). A locus pair was considered a true positive if any of the pairwise combinations of the genes assigned to the two loci were found to interact in the DKO dataset. Receiver operating characteristic (ROC) and precision-recall (PR) measures created using the ’precrec’ R package [69].

### Expression quantitative trait data

For the first real data benchmark, a dataset from an RM×BY yeast cross was used, comprising of genotype information along with gene expression measurements for 112 segregants [54]. We used the same set of unique loci as for the simulations. Because the methods would be evaluated based on the recovery of interactions for growth phenotypes (DKO dataset), 1050 transcripts were selected that code for essential genes according to the Saccharomyces Genome Deletion Project (www-sequence.stanford.edu/group/yeast_deletion_project/deletions3.html, [70]). This was done based on the assumption that genetic loci regulating the expression of essential genes are more likely to also affect growth than loci regulating non-essential genes. The expression values were scaled and centered to mean zero and equal variance.

The population structure covariates were created as follows: first, the relatedness, or kinship, between strains was estimated using the R package ’emma’ [71]. The first six eigenvectors, explaining 60% of genotypic variance, were used as representatives for the population structure. For the benchmark, the loci were mapped to nearby genes according to the following procedure: for each locus, all variants that were in LD (absolute correlation coefficient above 0.9) and that were less than 50 kilobasepairs (kbp) apart, were grouped. The locus region was then defined as starting one basepair (bp) after the previous locus and ending one bp before the next locus, or at the start or end of each chromosome for the first and last locus, respectively. All genes that overlapped this variant region according to the ENSEMBL annotation [72] were then assigned to the locus. To evaluate the performance of the methods at identifying interactions, the results for the different transcripts were grouped by taking the mean *p*-value across the transcripts for each locus pair.

To evaluate the interactions that were detected by each method exclusively, we did the following: for each transcript, and for each considered method (i.e., the three RF-based approaches, the exhaustive LM, the t-test 2step one, the LASSO 2step one, and the RF 2-step one), we extracted the interactions with the lowest *p*-value that was found by this method (*p*-value non-NA and in the lower quartile of *p*-values), but not by any of the other considered approaches (*p*-value NA or bigger than its 25% quartile of *p*-values), and which were also considered to be true positives according to the DKO dataset. As a reference, we also extracted the interaction with the lowest mean *p*-value (over all considered methods) that was found by at least 5 of the 7 methods (non-NA and among the top 500 *p*-values of each method).

### Growth trait dataset

For the evaluation of the methods on more complex traits, another yeast cross dataset was selected [60]. The dataset encompassed genotype information for 720 distinct segregants of a cross between the strains BY4742 and SK1 as well as measurements for several growth-related phenotypes. We selected the phenotypes ’growth rate on agar’ and ’resistance to cantharidin’ (a monogenic trait) as positive and negative control phenotypes, respectively. The phenotype values were scaled and centered to mean zero and equal variances. The original dataset encompassed genotype information for 65,250 genetic variants. Variants in high LD were grouped in the following way: For each chromosome, a hierarchical clustering of the variants was performed, using absolute Pearson correlation values subtracted from one as distance measures. The clustering tree was then cut at a height of 0.02. In the majority of cases, the clusters grouped variants that were located consecutively on the chromosome. For the resulting 2,827 loci genome-wide, a ’consensus locus’ was created for each cluster by using the majority vote (rounded mean of alleles encoded as 0 or 1) of the variants in each group. If the mean was between 0.4 and 0.6 the genotype was treated as unknown. The resulting minor allele frequencies ranged from 0.3 to 0.7, and there were no loci with more than 11% missing genotypes.

The population structure was estimated in the same way as for the eQTL dataset. Five principal components, explaining 55% of genotypic variance, were used as representatives for the population structure. The original 65,250 variants were mapped to genes in the following way: each (original, non-grouped) variant was defined as starting one bp after the previous variant and ending one bp before the next variant, or the start or end of the chromosome for its first and last variant, respectively. If there was a gene overlapping this region according to the ENSEMBL annotation [72], it was assigned to the variant. Finally, the ’consensus locus’ of each variant group was assigned all genes that were assigned to the variants in its locus.

## Supporting information

**S1 Fig. Different models of epistasis for a quantitative trait.** Depicted are examples of different models of epistasis between two hypothetical loci A and B. On the left side, additive effects are shown. The phenotypes expected under additivity are indicated by the stacked bars. All types of epistasis are also applicable to negative trait values. (A) Examples for positive (enhancing, or synergistic), negative (alleviating, or antagonistic), and reciprocal sign epistasis. (B) Example for pure XOR epistasis, where the marginal effects of two alleles are nullified if they occur together. (C) Example for pure AND epistasis, where two alleles only show an effect if they occur together. XOR and AND-type epistasis are not necessarily exclusive from the epistasis types shown in (A), but rather special cases of them.

**S2 Fig. Schematic representation of the Random Forest algorithm.** For each tree, a bootstrap sample of the individuals is taken. The locus that explains the most of the variance of the trait value is used to split the data into two groups. Here, the selected locus is represented as a dot at the splitting point in the color corresponding to the locus. The two subgroups are then again split on another locus, following the same procedure. At each splitting point, only a random subset of available loci is evaluated (represented by the dice).

**S3 Fig. Sensitivity of methods based on simulations of pure two- and three-way AND epistasis.** (A) Pure AND epistasis with small effect size (simulation scenario EA2); (B) Pure three-way AND-type interaction with bigger effect size (scenario EA3); (C) Pure three-way AND epistasis with small effect size (EA4).

**S4 Fig. Sensitivity of methods for scenarios with multiple interaction and marginal effects.** (A) to (D) show simulation scenarios M5, M7, M6, and M8. Each scenario encompasses two interactions and two marginal effects. Scenarios where the interacting loci additionally had a marginal effect (A and C) are compared to cases where the loci with marginal effects were distinct from the interacting loci (B and D). The interactions in (A) and (B) have a bigger effect size than the interactions in (C) and (D).

**S5 Fig. Sensitivity of methods for AND epistasis with and without marginal effects of the interacting loci.** Simulation scenarios M10 and M11 are depicted in (A) and (B), respectively. In (A), the marginal effect is on one of the interacting loci, while the marginal effect in (B) is on a distinct locus.

**S6 Fig. Influence of increasing number of marginal effects on interaction detection sensitivity.** (A) to (D) correspond to scenarios M1 to M4, respectively. The simulations encompass one AND-type interaction each, as well as marginal effects for one to four separate loci, respectively.

**S7 Fig. Sensitivity of methods based on simulations of XOR epistasis.** (A) Pure XOR epistasis with effect size 0.5 (simulation scenario EX2); (B) XOR epistasis with effect size 1 with additional marginal effects (size 1) on both interacting loci (scenario M9). The addition of these marginal effects did not improve the recovery of XOR epistasis.

**S8 Fig. Features of the interactions detected exclusively by each method.** (A) The interactions that were detected by each method, but not the others, and which were also considered to be true positives according to the DKO dataset, were extracted. These were then classified as either being XOR, or AND epistasis. The bar heights differ between methods because there was not always an interaction that was exclusively detected by each method for each transcript. (B) Same as (A), but interactions were classified based on whether the interacting loci had marginal effects, as indicated by a *t*-test *p*-value smaller than 0.05. (C) Here, it is indicated whether the strongest marginal effect (*t*-test) for each transcript was also one of the interacting loci for each method-exclusive interaction, or, in other words, whether there was a big marginal effect on a locus besides the interacting loci. (D) Boxplot of the number of locus pairs that were tested per transcript by each method, reflecting the nature of the different methods.

**S9 Fig. Meaningfulness of interactions for cantharidin resistance.** Locus pairs corresponding to gene pairs that were found to interact in the dataset were treated as true positives. Since cantharidin resistance is a monogenic trait in yeast, interactions were not expected to be meaningful, i.e., show an overlap with the DKO dataset. (A) precision recall (PR) curve. (B) Area under receiver operating characteristic (AUROC), with the values expected under random assignment subtracted.

